# Impact of *Capsicum frutescens (L.)* fruit ethanol extract on ibuprofen-induced gastric ulcers: A Wistar rat model

**DOI:** 10.1101/2025.06.07.658450

**Authors:** Joy Suubo, Herbert Izo Ninsiima, Miriam Nansunga, Edward Jessero, Etah Nkanu

## Abstract

Gastric ulcers are due to an imbalance between the protective factors like mucus and bicarbonate and offensive factors like gastric hydrochloric acid, infection and pepsin, on the other hand and medicines like ibuprofen. Despite the available conventional therapy, the mortality rate due to gastric ulcers across the globe is 11% with a range of 3-23%. *Capsicum frutescens* (*C. frutescens)* fruit which is consumed in most communities as a spice has a number of pharmacological benefits which include being an anti-ulcer, antimicrobial, anti-diabetic and anticancer activity, among other uses. This study was therefore carried out to determine the effect of the ethanol fruit extract of *Capsicum frutescens* on the gastric acid production determinants in ibuprofen induced gastric ulcers in Wistar rats.

Fifteen male Wistar rats were randomly assigned to five groups of three rats each. Group one (control) received normal saline, Other groups received ibuprofen for five days followed by normal saline, or misoprostol^R^, or *Capsicum frutescens* (107.5mg/kg body weight) for fourteen days. Ulcer index, H^+^/K^+^ ATPase enzyme activity, gastrin secretion and parietal cell histology were determined.

Oral administration of Ibuprofen induced gastric ulcers. *Capsicum frutescens* reduced the ulcer index. The H^+^/K^+^ ATPase enzyme activity was significantly decreased (p < 0.01) as well as concentration of gastrin (p < 0.01) and significant (p < 0.01) decrease in the mean number of parietal cells.

It is concluded that *Capsicum frutescens* has a therapeutic property for ulcer healing and does this through reduction of; H^+^/K^+^ ATPase enzyme activity, gastrin release and number of parietal cells in the stomach.

## Introduction

Gastric ulcers are a prevalent gastrointestinal disorder, affecting populations worldwide. In Western countries, the prevalence is approximately 2.4%, with an annual incidence ranging from 0.10% to 1.19%[1]. In contrast, regions such as Mainland China report a higher prevalence of 6.07%, with 22.5% of patients presenting with gastrointestinal symptoms diagnosed with gastric ulcers [2]. In Uganda, a prevalence of 2.9% was reported in 2015[3], with a significant disease burden, particularly in underserved populations with limited access to healthcare. The mortality rate in Uganda due to gastric ulcers stands at 2.81 per 100,000 individuals, underscoring the need for effective and accessible treatment options[1].

Gastric ulceration is characterized by mucosal damage extending to or beyond the submucosal layer of the gastric wall [4]Its pathophysiology involves an imbalance between protective factors, such as mucus and bicarbonate, and offensive factors, including gastric acid, Helicobacter pylori (H. pylori) infection, and pepsin [5]. Various risk factors contribute to gastric ulcer development, including genetic predisposition, excessive alcohol consumption, smoking, frequent starvation, stress, and prolonged use of non-steroidal anti-inflammatory drugs (NSAIDs)[6]. Oxidative stress, mediated by reactive oxygen species, exacerbates mucosal injury through lipid peroxidation, leading to cell membrane disruption and subsequent mucosal damage[7].

Current pharmacological interventions for gastric ulcers include antimicrobial agents, proton pump inhibitors (PPIs), histamine receptor antagonists (H2 receptor antagonists), antacids, prostaglandin analogs, and mucosal protective agents [8]. While effective, these treatments are costly and associated with adverse effects, including gastrointestinal disturbances, pancreatitis, hormonal imbalances, and contraindications in pregnancy [9] [10] [8]. Given these limitations, there is growing interest in alternative therapeutic strategies utilizing natural compounds with gastroprotective properties.

Capsicum frutescens (C. frutescens), commonly consumed as a spice, has demonstrated pharmacological benefits, including anti-ulcerative, antimicrobial, anti-diabetic, and anti-cancer activities [11]. *C. frutescens* has been shown to have an ulcer healing and gastric mucosa protective effect [12]. Studies suggest that prolonged use of C. frutescens for about two weeks enhances mucus and bicarbonate secretion while promoting prostaglandin E production, which inhibits gastric acid secretion [13–15]. Adequate gastric mucus, bicarbonate and prostaglandin E production are among the gastric mucosa protective factors [16]. However, short term use of *C. frutescens,* for one to three days or less, aggravates gastric ulcers through initiation of inflammatory reactions which leads to irritation of the gastric mucosa and pain. This was revealed by Jolayemi and Ojewole in the study they carried out in 2013 in Australia to investigate the comparative anti-inflammatory properties of capsaicin and ethyl-acetate extract of *C. frutescens* in rats [17]. Inflammation mobilizes defense mechanisms initiated by capsaicin which is a counter-irritant and in the long run counters the acid production with prolonged application of *C. Frutescens* [18]).

[19] showed that *C. frutescens* contains vitamin C, vitamin B_6_, vitamin B_3_, vitamin E, potassium, manganese, carotenoids [19] and flavonoids which are anti-oxidants among other biologic functions [20]) which could be one of the properties responsible for the gastric proactive mechanisms. According to [17], its fruit contains capsaicin (8-methyl-*N*-vanillyl-6-nonenamide) plus many other chemicals which have a number of similar branched and straight chain alkyl vanillylamides, generally known as capsaicinoids, and this is their main chemical constituent. Peverly, Salamova and Hites (2015) documented that the major capsaicinoids in *C. frutescens* are capsaicin which contributes 48.6% of the capsaicinoids, then 6, 7-dihydrocapsaicin. The minor capsaicinoids that are present are nordihydrocapsaicin (7.4%), homodihydrocapsaicin (2%), and homocapsaicin (2%). The seeds have the steroidal glycosides capsicoside A, B, C and D, and furostanol.

This study elucidates the mechanism by which C. frutescens modulates gastric acid production in ibuprofen-induced gastric ulcers in Wistar rats. Specifically, the study examines the effects of C. frutescens ethanolic extract on H+/K+ ATPase enzyme secretion, gastrin hormone levels, and parietal cell density.

## Materials and Methods

### Study Design

This experimental study investigated the effect of *Capsicum frutescens* fruit ethanol extract on gastric acid production determinants in ibuprofen-induced gastric ulcers in Wistar rats. The study involved 15 male Wistar rats, which were randomly grouped into five groups of three each. The rats received different treatments based on a previously established dosage from [21], demonstrating the potential of *C. frutescens* in reducing gastric acid production.

### Study Population

#### Animal Selection and Housing

Fifteen male Wistar rats (6-8 weeks old) were obtained from the Pharmacology Laboratory of Kampala International University-Western Campus. They were housed in cages with a glass bottom and stainless-steel mesh walls for aeration, provided with a 12-hour light/dark cycle, and fed standard rat chow and water *ad libitum*. Prior to the experiment, the rats were weighed to ensure accurate dosing.

#### Treatment Groups

The rats were randomly assigned to five groups, each receiving different treatments:

#### Sampling Techniques

A simple random sampling technique was used to allocate the rats to their respective groups. Each rat was assigned a unique number, and selection was done randomly using Microsoft Excel 2010.

#### Sample Size Determination

The sample size was calculated using the formula: n=DFK+1n = \frac{DF}{K+1} where *n* is the number of rats per group, *DF* is degrees of freedom (10 to 20), and *K* is the number of groups. The calculation yielded 2 to 3 rats per group, and the final sample size was set at 15 rats (3 per group) following previous studies [21].

### Experimental Procedures

#### Plant Collection and Identification

Approximately 400g of ripe *C. frutescens* fruits were collected from Rwemirokora village, Bushenyi District, Uganda. The fruits were shade-dried for 14 days, and botanical identification was done at Makerere University Herbarium, receiving voucher specimen number Mak 1911-JS076.

#### Extraction of *C. frutescens* Fruit

The dried fruits were powdered and extracted using 70% ethanol via cold maceration. The extract was filtered, oven-dried at 50°C, weighed, and stored at 4°C.

#### Induction and Confirmation of Gastric Ulcers

Gastric ulcers were induced using ibuprofen (40mg/kg body weight) administered oro-oesophageally for five consecutive days. The development of ulcers was confirmed based on clinical signs (abdominal scratching and biting) and histological examination in a pilot study.

#### Preparation and Administration of Extract and Drugs

After obtaining *C. frutescens* fruit extract by the method described in 3.6.2, a pre-weighed mass of the extract was suspended in normal saline. After thorough stirring, the drug was administered according to the formula;

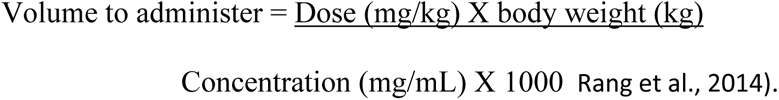

### Data Collection and Analysis

#### H+/K+ ATPase Enzyme Activity

Stomachs were excised post-euthanasia and processed to isolate gastric microsomes. ATPase activity was measured spectrophotometrically at 700nm by quantifying inorganic phosphate release using a molybdenum blue assay.

#### Gastrin Hormone Quantification

Plasma gastrin levels were measured using an ELISA assay after fasting the rats overnight. Blood was collected via cardiac puncture, centrifuged to obtain plasma, and analyzed at Makerere University’s Biochemistry Department.

#### Histological Analysis of Parietal Cells

Gastric tissue samples were fixed in 10% formal saline, processed for histology, and stained with Hematoxylin and Eosin (H&E). Parietal cell count was determined under a light microscope at 40x magnification based on characteristic histological features.

#### Ulcer induction

Ibuprofen tablets (Lincoln Pharmaceuticals limited) were crushed to powder and suspended in normal saline for easy administration. Administration started after 24 hours of starvation of the rats. The dose rate was 40mg/kg body weight once daily. Four hours post administration of ibuprofen tablets, there were no clinical signs of gastric ulcers and when the rats were opened up surgically, no ulcers had been induced. After three days of consecutive administration of ibuprofen tablets (Lincoln Pharmaceuticals limited) at a dose rate of 40mg/kg body weight once daily, the mucus on the gastric mucosa was depleted but no ulcers had been induced yet. Five days post consecutive day administration of ibuprofen tablets (Lincoln Pharmaceuticals limited), the rats were showing clinical signs of biting and scratching around the abdominal area, and when opened up surgically, gastric ulcers had been induced as shown in Fig 1.

**FIGURE 1:**
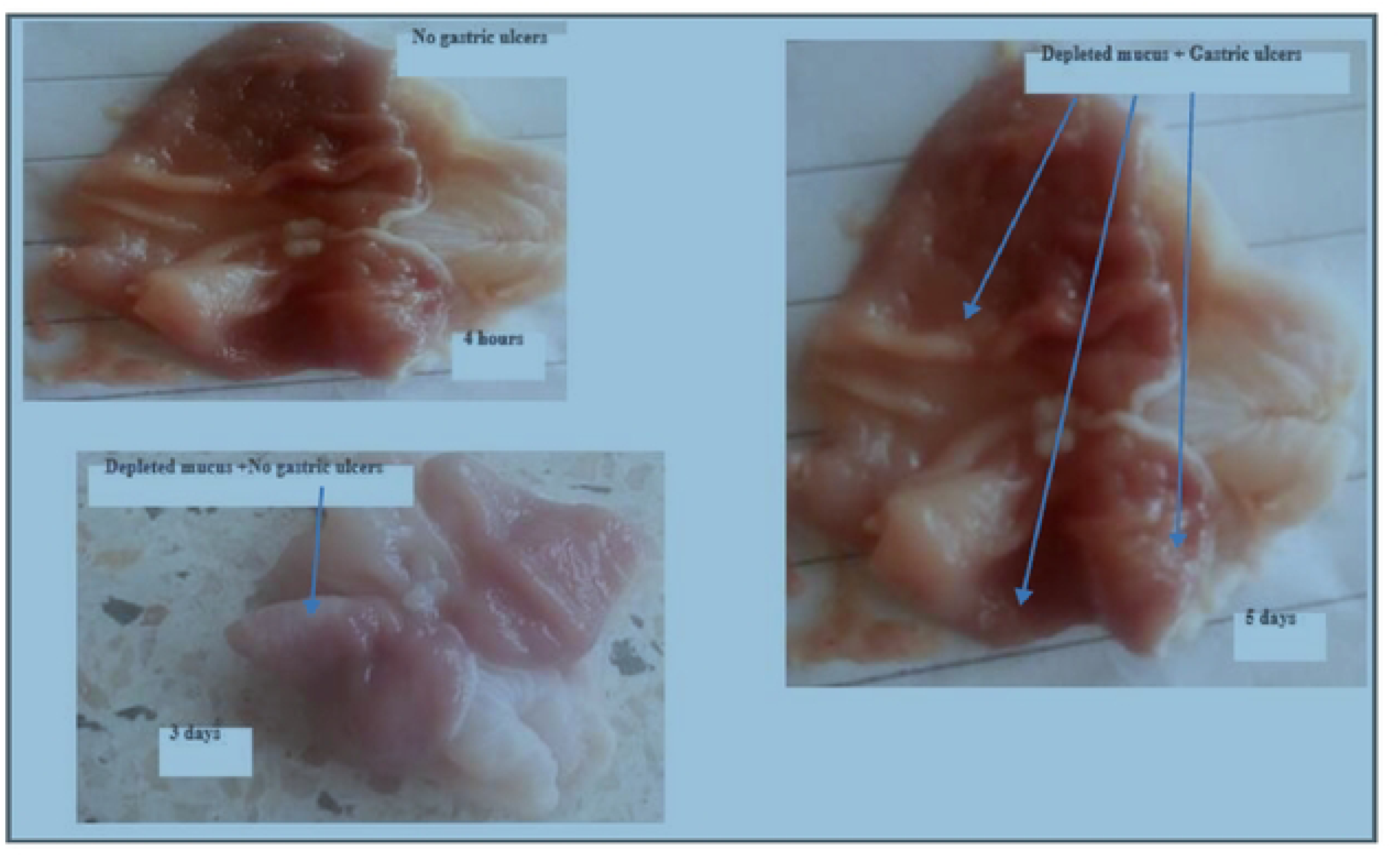
GASTRIC ACID INDUCTION

## RESULTS

### Ulcer index

Of the 12 rats that were induced with gastric ulcers, 3 (25%) of those that received normal saline still had ulcers at the end of the 14 days of treatment while 9 (75%), that received misoprostol and the different concentrations of *Capsicum frutescens* extract had no gastric ulcers. The ulcer indices were as shown in table 2.

**Table 1:** Study groups and treatment.

| Group | Treatment |
| --- | --- |
| Normal Control | No ulcers + Normal saline (2ml/kg body weight) |
| Negative Control | Gastric ulcers + Normal saline (2ml/kg body weight) |
| Positive Control | Gastric ulcers + Misoprostol (100µg/kg body weight, once daily for 14 days) |
| Experimental Group I | Gastric ulcers + *C. frutescens* fruit extract (107.5mg/ml, once daily for 14 days) |
| Experimental Group II | Gastric ulcers + *C. frutescens* fruit extract (215mg/ml, once daily for 14 days) |

**Table 2:** Showing the treatment given to the different cases and their outcome on ulcer indices

| Treatment group | Treatment | Dose rate |
| --- | --- | --- |
| Ibuprofen + normal saline | Normal saline | 2ml/kg bw |
| Ibuprofen+ misoprostol | Misoprostol | 100µg/kg bw |
| Ibuprofen+ *C. frutescens* | *C. frutescens* | 107.50 mg/kg bw |
| Ibuprofen+ *C. frutescens* | *C. frutescens* | 215 mg/kg bw |

With a mean ulcer index of 1.13 cm, rats in the control group who were given normal saline (2 ml/kg body weight) showed persistent ulcers with ulcer indices which varied from 1.0 to 1.3 cm. Rats given misoprostol (100 µg/kg body weight) on the other hand displayed full ulcer healing, with a mean ulcer index of 0.0 cm, indicating that misoprostol was very successful in ending ulcers. Likewise, all rats treated with extract from Capsicum frutescens at 107.50 mg/ml and 215 mg/ml showed full ulcer healing, with an average ulcer index of 0.0 cm.

Overall, the findings show that whereas normal saline had no therapeutic effect on gastric ulcers, C. frutescens extract at the tested doses considerably improved ulcer healing and was comparable to the common anti-ulcer medication, misoprostol.

#### The H^+^/K^+^ ATPase enzyme activity analysis

Administration of Capsicum frutescens fruit extract significantly reduced the H/K ATPase activity as shown in Fig 2.

**Fig 2:**
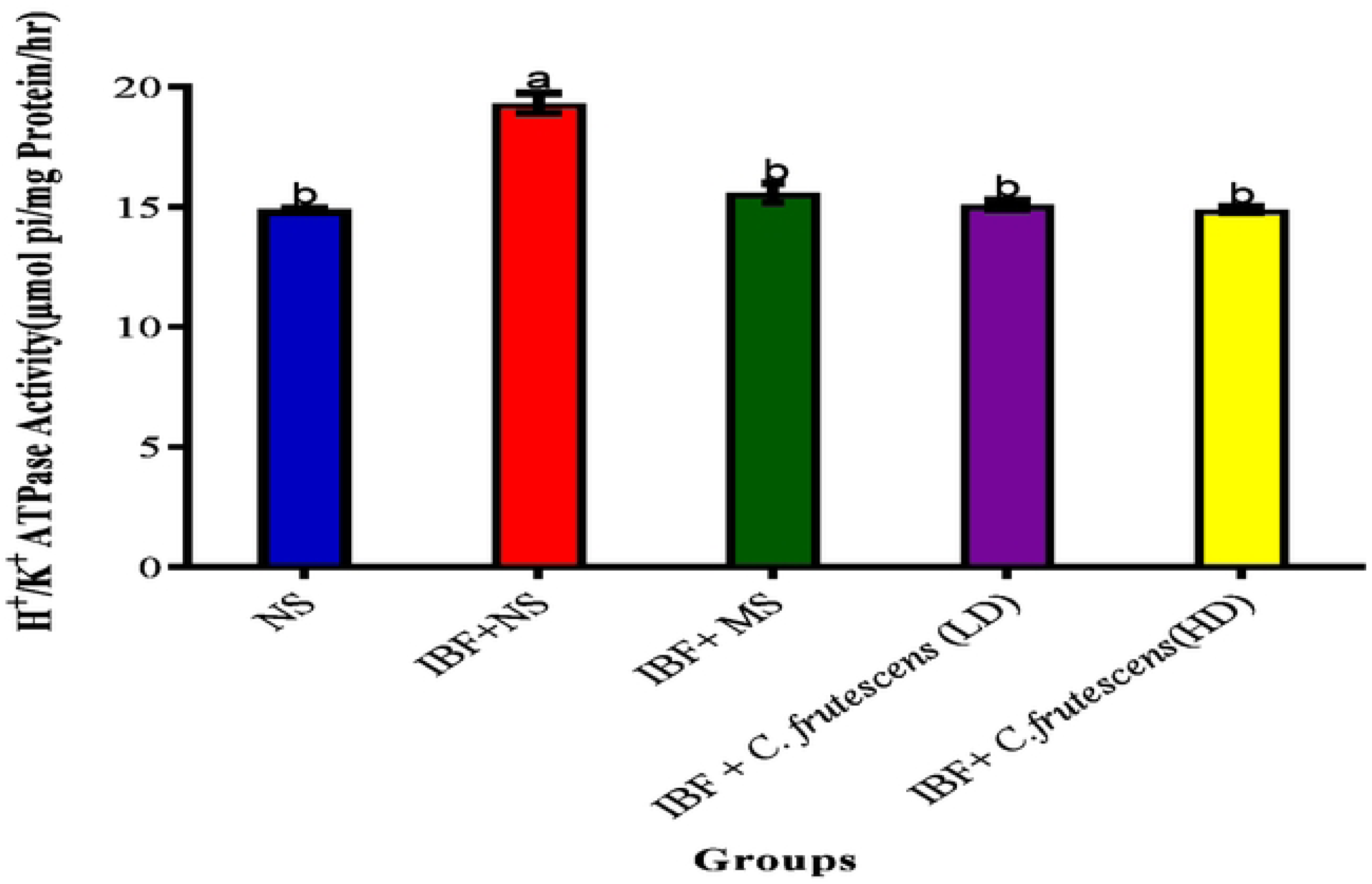
Showing variation of the activity of *H+JK+* ATPase enzyme across the groups. b=p<:0.05 vs Control and other experimental groups. a=NS when compared to control and between groups. Values are expressed as Mean±**SEM**

The group that received normal saline treatment (control) for stomach ulcers showed the highest enzyme activity that was significantly different from other treatment groups. On the other hand, misoprostol-treated rats exhibited a significant decrease in H⁺/K⁺ ATPase activity, indicating that misoprostol efficiently suppresses stomach acid output and aids in ulcer healing. Similar to the group treated with misoprostol, both doses of Capsicum frutescens (A and B) produced decreased levels of enzyme activity. There was no significant difference between those treated with misoprostol and Capsicum frutescensis.

#### Quantity of gastrin hormone analysis

Ibuprofen administration caused an increase in gastrin hormone secretion. Capsicum frutescens administration (A and B) significantly reduced gastrin hormone levels compared to the ulcer control, with values approaching those of the non-ulcerated group as shown in Fig 2. similarly, rats treated with misoprostol had lower gastrin levels than the ulcer control group

#### Analysis of number of parietal cells

Capsicum frutescens fruit extract (A and B) significantly resuced the number of parietal cells as shown in Fig 3 and 4. The gastric ulcer group treated with normal saline had the greatest number of parietal cells, as shown by statistical annotation (b), implying that ibuprufen is associated with an increased parietal cell population, which may contribute to excessive gastric acid secretion and ulcer persistence.

**Fig 3:**
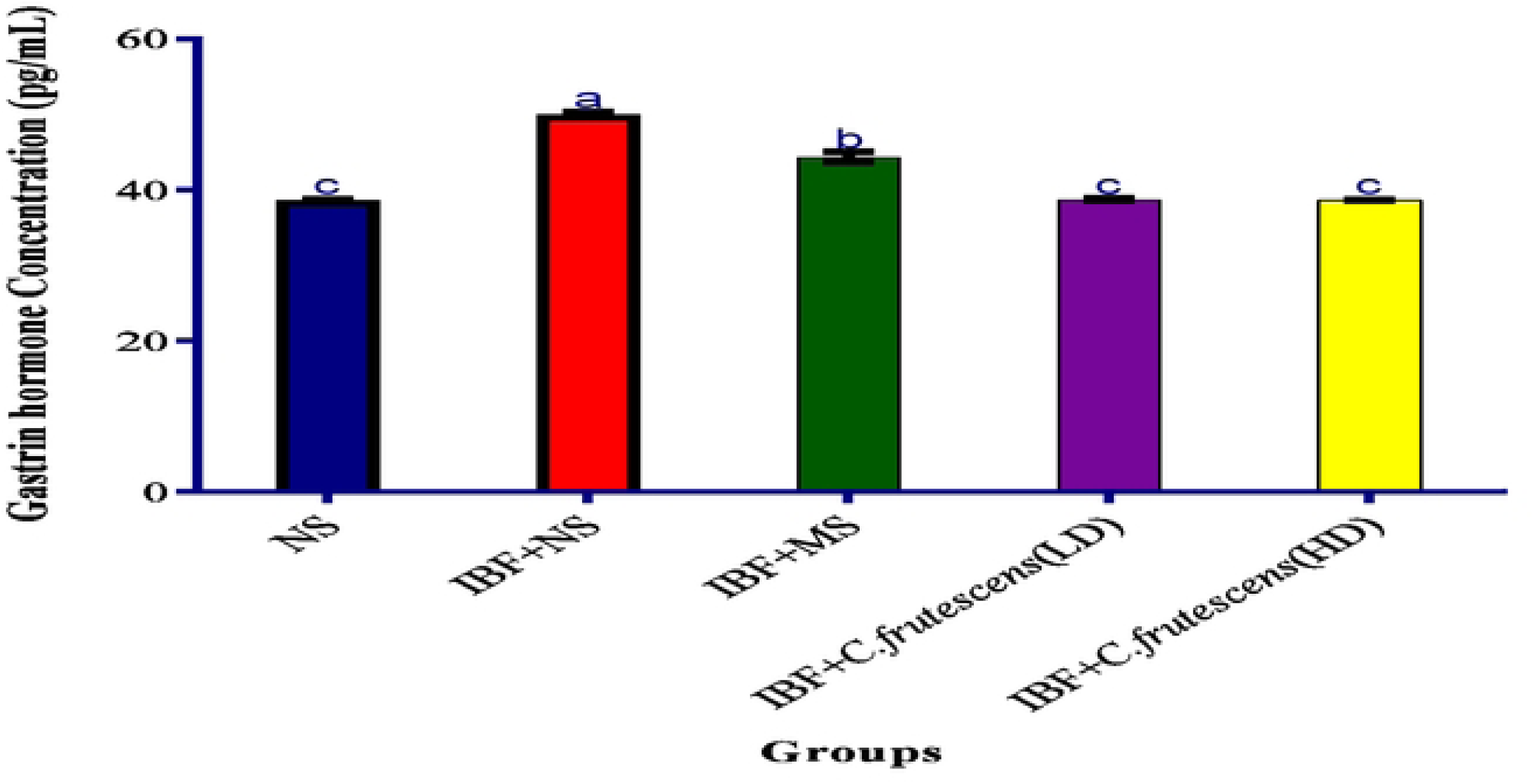
Showing the variation of the quantity of gastrin hormone across the treatment groups. Same letters indicate no statistical difference and different letters indicate statistical difference. B=p<0.05 vs Control and other experimental groups. a=NS when compared to control and between groups. c=p<0.05 vs control and other experimental groups. Values are expressed as Mean ±SEM.

**Fig 4:**
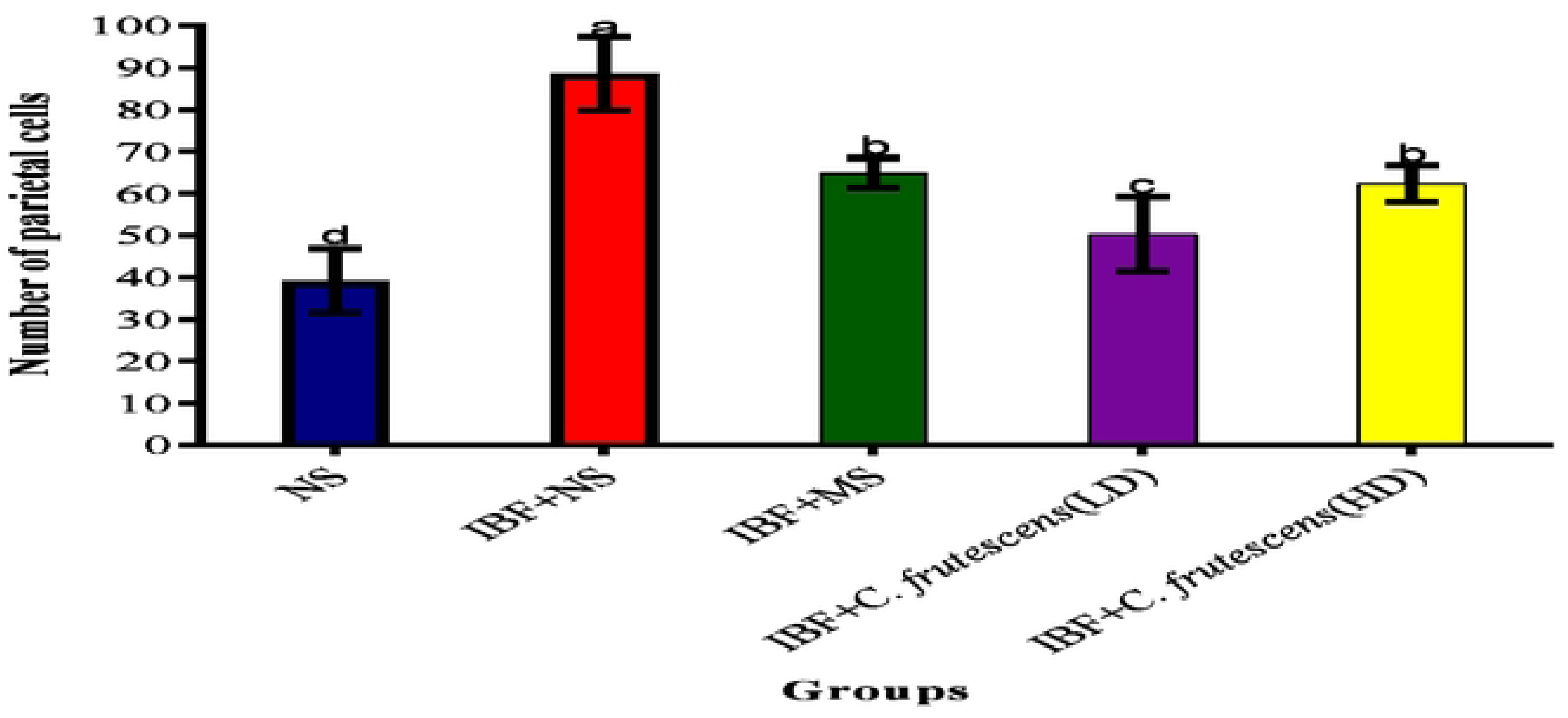
Showing variation of mean number of parietal cells across the treatment groups. Same letters indicate no statistical differences while different letters show statistical differences between the groups. b=p<0.05 vs Control and other experimental groups. a=NS when compared to control and between groups. c=p<0.05 vs control and other experimental groups. Values an: expressed as Mean ±SEM.

**Fig 5.**
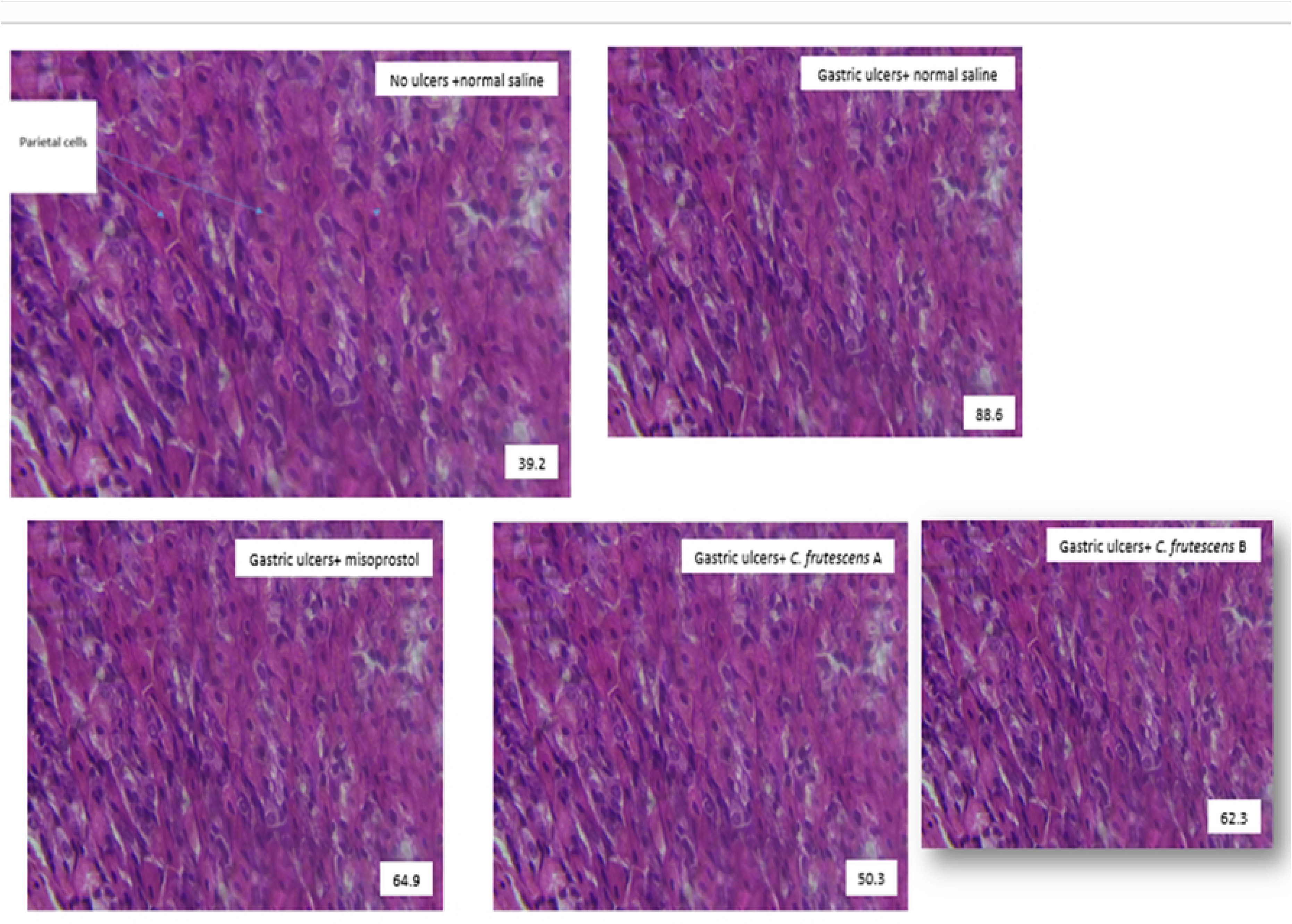
Effect on parietal cell histology.

When compared to the ulcer control group, the number of parietal cells decreased as a result of treatment with capsicum frutescens (A and B), with the impact being more noticeable in C. frutescens B. Both dosages decreased the number of parietal cells, but the larger dose had a greater effect on limiting the cells that secreted acid, according to the statistical markers (a for C. frutescens A and c for C. frutescens B**). Analogously, misoprostol administration led to fewer parietal cells, as seen in annotation (c).

The non-ulcerated control group contained a moderate number of parietal cells, indicating normal stomach physiology.

## Discussion

This study reports on the effect of *Capsicum frutescens (L.)* fruit ethanol extract on gastric acid production determinants in ibuprofen-induced gastric ulcers in Wister rats.

In this study ibuprofen at 40mg/kg weight per day for 5 days induced gastric ulceration, increased the H+/K+ ATPase activity, increased gastrin secretion and number of parietal cells. Other studies have shown similar mechanisms which could be due to its irritant effect on the stomach wall and inhibition of prostaglandin secretion[23] [24].

This study observed the positive curative potentials of *C. frutescens* via the reduction of ulcer index, H^+^/K^+^ ATPase activity, reduced gastrin concentration as well as reduced number of parietal cells, which is a major source of hydrochloric acid [25].

There were no registered gastric ulcers observed in rats which receive *C. frutescens* extract as shown by an ulcer index of 0 compared to a positive control group. This shows the curative potential of C. frutescens as shown by other previous studies.[25]. This could be due to the different mechanisms like increased mucus secretion and reduced hydrochloric acid secretion from parietal cells that enhances ulcer healing. Previous research have shown that *C. frutescens* can heal gastric ulcers through increasing production of gastric mucus [21], prostaglandins which reduces gastric hydrochloric acid production while increasing secretion of alkaline thick mucus and bicarbonate, hence aiding in protection of the gastric mucosa from direct contact with acid [26]

The present study explored the different mechanisms of gastric ulcers healing using *C. frutescens* extract including H+/K+ ATPase, gastrin secretion and parietal cell growth.

*C. frutescens* extract significantly reduced the H+/K+ ATPase activity which is responsible for the increased hydrochloric acid production [27]. H+/K+ ATPase upregulation is implicated in ulcer disease as evidenced by the significant increase observed in the ulcer control group. The plant extract may be a viable natural substitute for controlling excessive acid output in gastric ulcer settings, as indicated by the comparable decreases seen in the misoprostol and C. frutescens-treated groups. The therapeutic property of *C. frutescens* shown in this work may be attributed to the phytochemical components of *C. frutescens* such as carotenoids, vitamin C, B3, B6 and E as well as manganese, flavonoids and potassium, which are anti-oxidants hence promote healing of ulcers [28]. Therefore, the possible inhibition of H^+^/K^+^ ATPase enzyme activity and gastrin secretion by *C. frutescens* amongst other mechanisms for healing gastric ulcers cannot be over emphasized. It is therefore also suggestive that *C. frutescens* may be acting through inhibition of the proton pump by binding to the H^+^/K^+^ ATPase pump on the parietal cells. Parietal cells in the stomach are known for their hydrochloric acid production with a significant role of protecting against bacterial infection. Drastically increasing the number of these cells poses danger to the mucosal walls and may promote excoriation. Interestingly, the remarkable increase of number of parietal cells caused by ibuprofen in this study was down regulated by *C. frutescens*. This correlates with a reduction in amount of secretion of hydrochloric acid as reported by [7]. The significant increase in parietal cells in this group aligns with the previously observed elevated H⁺/K⁺ ATPase activity and gastrin levels, reinforcing the role of parietal cell hyperplasia in ulcer pathology.

## Ethical considerations

Care for the animals was done according to the National Institute of Health, 2006 guidelines for use and care of laboratory animals in research teaching. The rats were fed standard rat meal and water ad lib and they were kept in cages made of a glass bottom and top and stainless-steel mesh wall for aeration. Light was availed twelve hours a day. When carrying out the experiment, all treatments, that is ibuprofen, misoprostol and *C. frutescens* fruit ethanol extract were administered in doses already proven not to be harmful and documented. When confirming induction of the gastric ulcers, a combination of ketamine and xylazine were administered as anaesthetics to minimise pain to the animals. During euthanasia, thiopental sodium was administered so that the animals die humanely. In order to protect the environment, the carcasses were incinerated immediately after death. Before starting the study, permission was sought from the department of physiology KIU-WC, faculty of biomedical sciences KIU-WC, Postgraduate Studies and Research Directorate (PGSRD) KIU-WC and KIU Research Ethics Committee (REC) and ethical approval number is Nr.UG-REC-023/201943.

## Conclusion

Based on our findings in this research work, we conclude that *Capsicum frutescens* has a therapeutic property for ulcer healing and does this through reduction of H^+^/K^+^ ATPase enzyme activity, gastrin release and number of parietal cells in the stomach.

## Acknowledgement

I thank Dr. Justin Atiang Beshel, Wilber Ntoni, Christopher Nandala and Dr. Muhammudu Kalange for their tireless contributions in this work.

I also thank my mother Mrs Tabitha Baleeta, sisters (Harriet Abenakyo and Mary Babirye) and brothers (Fred Baleeta, Daniel Idibya, Henry Idibya and Emmanuel Kumbuga) for the support and encouragement given me during this research.

I thank Mr Ambrose Mukisa, the Chief laboratory technician, Biochemistry department, Makerere University, for the assistance accorded to me when carrying out the experiments in the laboratory.

Finally, I really thank God, I give Him all the glory and honour.

---

**Platt 3:**
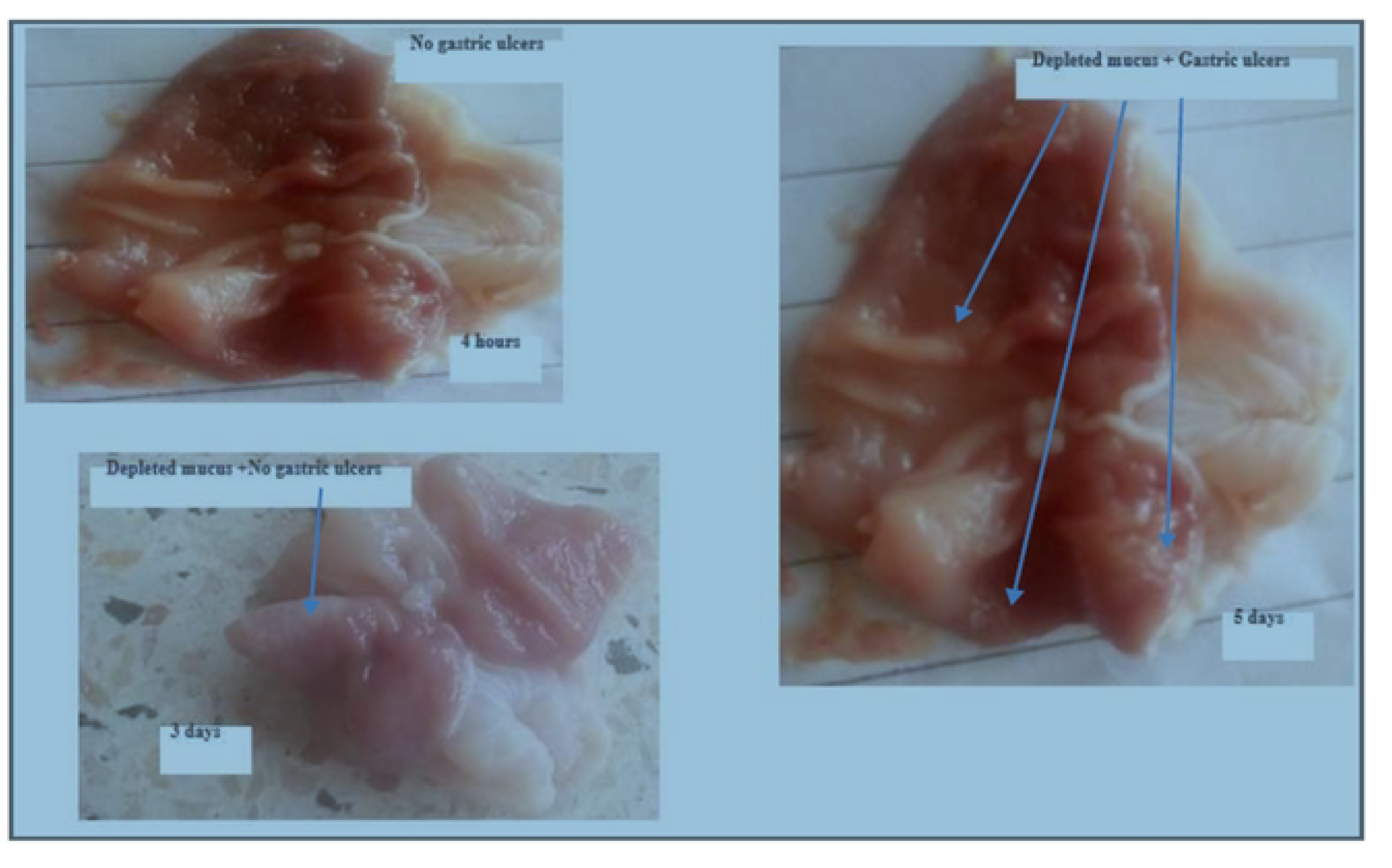
Pilot study results ou induction of gastric ulcers using ibuprofen.

